# Cortical, muscular, and kinetic activity underpinning attentional focus strategies during visuomotor control

**DOI:** 10.1101/2022.07.21.501019

**Authors:** J. V. V. Parr, G. Gallicchio, A. Canales-Johnson, L. Uiga, G. Wood

**Affiliations:** Department of Sport and Exercise Sciences, Manchester Metropolitan University Institute of Sport, Manchester, United Kingdom; School of Human and Behavioural Sciences, Bangor University, Bangor UK; Consciousness and Cognition Lab, Department of Psychology, University of Cambridge, Cambridge, United Kingdom; Viccerectoria de Investigacion y Posgrado, Universidad Catolica del Maule, Talca, Chile

**Keywords:** attentional focus, internal focus, corticomuscular coherence, frontal midline theta, EEG alpha activity, isometric force precision

## Abstract

Focusing internally on movement control or bodily sensations is frequently shown to disrupt the effectiveness and efficiency of motor control when compared to focusing externally on the outcome of movement. Whilst the behavioural consequences of these attentional strategies are well documented, it is unclear how they are explained at the corticomuscular level. The aim of the present study was to investigate how attentional focus strategies affect kinetic, cortical, muscular, and corticomuscular activity during an isometric force precision task. In a repeated measures design, we measured force, EEG and EMG activity from twenty-seven participants who performed 160 isometric contractions of the right hand whilst encouraged to adopt either an internal or external focus through a combination of instructions, secondary tasks, and self-report evaluations. Results indicated that focusing internally led to poorer force accuracy and steadiness compared to an external focus. An internal focus also increased muscle activity of the forearm flexor, increased EEG alpha activity across the parieto-occipital cortex, lowered frontal midline EEG theta activity, and lowered beta corticomuscular coherence between the forearm flexor and contralateral motor cortex. The results of this study provide a holistic understanding of how attentional focus strategies alter neuromuscular control during an isometric force precision task, paving the way for exploring how the behavioural consequences of attentional strategies can be explained at the corticomuscular levels across a wide range of motor tasks and contexts.

## 1. INTRODUCTION

Extensive research suggests that adopting an internal focus of attention, compared to an external focus of attention, is less effective for motor performance and learning (for reviews, see Wulf, 2007, 2013). An internal focus of attention occurs when an individual directs attentional resources inwards towards the control of movement or associated bodily sensations, whereas an external focus occurs when an individual allocates attentional resources towards the outcomes of the movement or the effects the movement has on the environment. The behavioural consequences of these strategies are well documented. For example, studies have shown that instructions promoting an internal focus can disrupt the accuracy (Bell & Hardy, 2009; Lohse et al., 2014), stability (Kim et al., 2017), and fluency of motor control (Kal et al., 2013).

A common mechanism purported to explain the disruptive effects of an internal focus is that it promotes “constrained action” consisting of elevated conscious control of movements through self-regulation, triggering the tendency to constrain the motor system and “freeze” the degrees of freedom (McNevin et al., 2003). By contrast, an external focus is proposed to place greater emphasis on the motor system’s self-organising capabilities, allowing for reflexive and automatic control. Interestingly, the constraining effect of an internal focus is supported by evidence that focusing internally elevates electromyographic (EMG) activity across antagonistic muscle pairs during isometric force production tasks, reflecting a “stiffening” strategy supported by co-contractions (Lohse et al., 2011; Lohse & Sherwood, 2012). An internal focus therefore appears to promote a state of diffuse muscular activity in a manner that is not supportive of efficiency and optimal task performance.

Although little is known about how the effects of an internal focus are explained at the cortical level, there is evidence that elevated EMG activity might be caused by a disruption to intracortical inhibition within the primary motor cortex (i.e., M1). Specifically, Kuhn et al. (2017a, 2017b) measured motor evoked potentials (MEPs) across several hand muscles in response to single-pulse transcranial magnetic stimulation to M1 during isometric contractions of the index finger. Their results showed that MEPs of the primary and surrounding muscles were greater when adopting an internal focus compared to an external focus, suggesting an internal focus diminished the activity of the inhibitory intracortical circuits within M1. Thus, focusing internally may disrupt the capacity to exert inhibitory control, leading to an overflow of activation at the cortex that becomes reciprocated at the muscular level.

There is also evidence that adopting an internal focus can have a significant effect on visual processing. For example, solving word generation tasks without the availability of task-relevant information (i.e., “in the mind’s eye”; internal focus) has been shown to elevate EEG alpha activity across posterior brain regions compared to when task-relevant visual information is available (i.e., the word; external focus; Ceh et al., 2020). Given that elevated EEG alpha activity is proposed to reflect regional inhibition in the cortex (Jensen & Mazaheri, 2010; Klimesch et al., 2007), these findings suggest that focusing internally may prevent visual inflow to shield ongoing internal processing (Benedek, 2018). These findings have recently been supported in the context of a single-limb balance task. Specifically, Sherman et al. (2021) found that focusing on knee angle position (i.e., internal focus) led to poorer balance performance and relatively greater EEG alpha activity across the visual and sensorimotor regions compared to when focusing on the position of a laser pointed towards a target (i.e., external focus). Interestingly, Sherman et al. also found that the internal focus condition decreased frontal midline theta activity compared to an external focus. Frontal midline theta is proposed to reflect the activity of the anterior cingulate cortex and is commonly measured as an index of top-down volitional attentional control to adjust behaviour in response to feedback error (Cavanagh et al., 2010; Cavanagh & Frank, 2014; Womelsdorf et al., 2010). Taken together, these findings suggest that focusing internally disrupts the detection and correction of somatosensory and visual information in a manner that impairs effective visuomotor control.

Whilst varying the attentional focus appears to drive changes in brain and muscle activity, it is unclear how these changes are related. Investigating functional interactions between the central and peripheral nervous systems may, therefore, provide new insights to the mechanisms underpinning the visuomotor response to attentional focus instructions. A promising technique to study cortical-muscular coupling during movement is corticomuscular coherence (CMC), a measure that evaluates the frequency-wise similarity between EEG and EMG signals (Mima & Hallett, 1999). CMC is typically observed during periods of isometric contraction (Kilner et al., 2003; Riddle & Baker, 2006) and reaches its peak in the beta frequency band (approx. 15 – 30 Hz) for EEG sites located over the primary sensorimotor cortex contralateral to the innervated limb (Witham et al., 2011). CMC is therefore proposed to reflect cortical control of motor unit firing via the direct corticospinal pathway (Mima & Hallett, 1999), building upon the presence of sensorimotor beta oscillations that are critical to maintaining a stable sensorimotor output (Bourguignon et al., 2019; Engel & Fries, 2010). Beta-range CMC increases after visuomotor skill learning (Perez et al., 2006), is higher during more precise force contractions (Kristeva et al., 2007), and is lower and less spatially localised in stroke patients (e.g., Larsen et al., 2017). Interestingly, beta band CMC also decreases significantly when attention is divided between a motor task, and another simultaneously performed task. Beta CMC may, therefore, reflect attention towards the task and serve to promote an effective and efficient corticospinal interaction (Kristeva et al., 2007), enhancing cortical control of muscle activity. As an internal focus is typically associated with poorer force control and elevated attentional demands, the typically detrimental effects of adopting an internal focus for motor performance may therefore be underpinned by a disruption of CMC.

The aim of the present study was to explore how varying the attentional focus alters force control, the activity of the cortex (via EEG), the muscles (via EMG), and their interaction (via CMC analyses) during a visually guided isometric handgrip task. Based on previous research, we hypothesised that internally focused instructions would encourage contractions that are less accurate, less stable, and supported by greater EMG activity of the forearm compared to external focused instructions. At the cortical level, we hypothesised that focusing internally compared to focusing externally would disengage visual processing (increased occipital alpha activity) and disrupt executive functions required for the detection and correction of visual feedback (decreased frontal midline theta activity). Finally, we hypothesised that an internal focus would impair beta CMC between the forearm muscles and the contralateral sensorimotor cortex compared to an external focus.

## 2. METHOD

### 2.1. Participants

Twenty-seven participants were recruited (17 males, 10 females, all right-handed, age M = 23.76 ± 4.34 years) based upon an a-priori sample size estimation performed using G*POWER software (version 3.1; Heinrich Heine University Dusseldorf, Dusseldorf, Germany). Whilst it is unclear how attentional focus strategies affect beta CMC, large effects have been observed when increasing attentional demands during motor tasks (Johnson et al., 2011; Johnson & Shinohara, 2012). If assuming a large effect (*d* = 0.80) with 80% power and an alpha level of *p* = .05 a sample size of at least 15 participants would have been required to compute the difference between two dependent means. However, we decided to increase our sample size to enable the detection of more subtle effects given the novelty of our study. Post-hoc sensitivity analyses therefore indicated that with 27 participants and an alpha level of *p* = .05, 80% power is achieved for medium effect sizes (*d* ≥ 0.56, ∼ *r* = .3). All participants were free from any known neurological/musculoskeletal disorders and were instructed to avoid stimulants (caffeine, alcohol, energy drink, etc.) 48 hours prior to testing. Ethical approval was granted by an institutional ethics committee and written informed consent was obtained from all participants.

### 2.2. Study protocol

#### Procedure

All participants attended the laboratory on one occasion for approximately two hours. Upon arrival, participants were debriefed about the study protocol before being sat comfortably at a desk facing a computer screen. Participants were then prepared for EEG and EMG data collection prior to performing two maximal voluntary contractions (MVC’s) separated by one minute of rest. The maximal force produced across these two attempts was used to personalise the target force output in the subsequent experimental task.

#### Experimental task

Participants were instructed to squeeze a dynamometer by contracting their right hand at 15 ± 5% of their MVC for five seconds. The force produced and target force boundaries (i.e., ± 5%) were displayed in real time on a monitor using Labchart 8.0 (Figure 1). Following a familiarisation block, participants completed 160 contractions, divided into 16 blocks of ten contractions. Each 5-second contraction was followed by a 5-second rest, and each block was separated by 30-seconds of rest to minimise muscular fatigue. The onset and offset of each 5-second contraction were indicated by auditory tones (10ms duration) controlled by PsychoPy (Peirce, 2007) and used as event triggers for the physiological and kinetic data. At the beginning of each block, participants were instructed to adopt either an external or internal attentional focus. The attentional conditions were presented in equal proportion (8 consecutive blocks per condition) and their order was permuted for each participant through PsychoPy. The attentional focus was manipulated through a combination of instructions, secondary tasks, and self-report evaluations.

**Figure 1.**
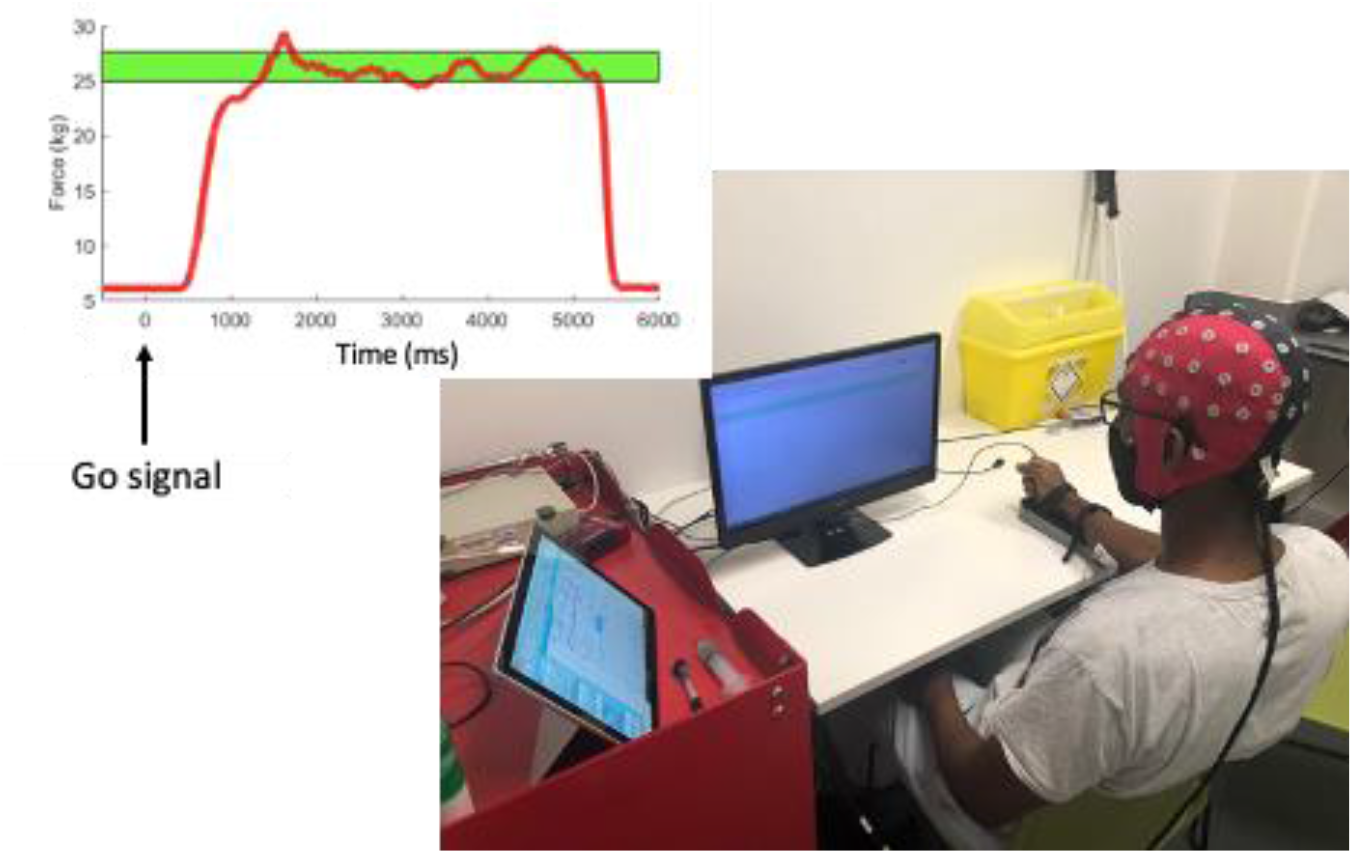
Visual representation of the experimental task set-up. The participant’s right hand is strapped in a cushion rig whilst wearing EEG and EMG equipment. Participants received ongoing visual feedback of force output on the monitor, trying to maintain the red force trace within a green boundary zone (depicted top left).

#### Attentional focus instructions

For the external focus condition, the following instructions were provided: “*For the next set of trials, carefully maintain your focus on the screen and the line being produced as you squeeze the dynamometer. Try and keep this line within the green boundary zone as accurately as possible*.” In addition, participants were asked to rate their accuracy on a scale from 1 (very inaccurate) to 5 (very accurate) at the end of each block. The secondary task required participants to verbally report whether the force line was “above” or “below” the mid-point of the green boundary zone upon the sound of a bell. This bell sounded 2.5 seconds into a randomly selected contraction during each block (controlled by Psychopy). Physiological and kinetic data recorded during trials containing a bell stimulus were removed from all analyses. Participants were informed the accuracy of their responses was being measured to assess their adherence to the task instructions.

For the internal focus condition, the following instructions were provided: “*For the next set of trials, carefully focus on the contraction of your forearm muscles as you squeeze the dynamometer. Try and keep the contracted force line within the green boundary zone as accurately as possible whilst applying an equal amount of force across your index and middle fingers.”* In addition, participants were asked to rate how equally they maintained the force across their index and middle fingers upon the dynamometer from 1 (not equally at all) to 5 (completely equally) at the end of each block. The secondary task required participants to verbally report whether the index or middle finger was applying the most force at the instance of the bell sound. Participants were told their ability to maintain equal force across the index and middle fingers would be assessed using the EMG electrodes.

#### Physiological and kinetic data

The EEG signals were recorded from 63 active shielded AgCl electrodes embedded in a stretchable fabric cap (eego sports, Ant Neuro, Hengelo, Netherlands) and positioned according to the extended 10-20 international system (Jurcak et al., 2007). Electrodes in sites CPz and AFz were used as reference and ground, respectively. Naison, Inion, and preauricular points were used as anatomical landmarks to position the EEG cap. Conductive gel for electrophysiological measurements was used (Signa gel, Parker), and impedance was kept below 20 kΩ. The EMG signals were recorded from two pairs of bipolar surface EMG electrodes (eego sports, Ant Neuro, Hengelo, Netherlands) placed on the skin to record activity from the flexor carpi radialis and extensor carpi radialis of the right forearm according to the guidelines set out by SENIAM (http://seniam.org/sensor_location.htm). The participant’s right forearm was then strapped to a cushioned rig positioned on the desk, where participants held a dynamometer connected to a PowerLab 4/25T (AD Instruments, Bella Vista, Australia) to record hand contraction force (in kilograms) via Labchart 8 software (ADinstruments). The force, EEG, and EMG signals were recorded at a sample rate of 1000 Hz and were synchronized through a square-wave trigger (i.e., time point zero in subsequent analyses) sent by a LabJack U3 device (LabJack Corporation, Lakewood, United States) at each auditory contraction prompt generated through a bespoke Psychopy programme.

### 2.3. Data processing

For EMG-specific analyses, signals were down-sampled (250 Hz), band-pass filtered (10 – 200 Hz), and notch filtered (48 – 52 Hz) prior to being cut into epochs ranging from -2 to +6 s relative to the onset auditory stimulus. For EEG and EEG-EMG analyses, signals were down-sampled (250 Hz) and band-pass filtered (1 – 45 Hz; finite impulse response) prior to being cut into epochs ranging from -2 to +6 s relative to the onset auditory stimulus. These epochs were visually inspected, and those showing large EEG contamination from muscular artefacts were discarded (from both EEG and EMG analyses). No bad EEG channels were identified. Independent component analysis (ICA) weights were obtained through the RunICA infomax algorithm (Jung et al., 1998) running on EEG signals. ICA weights that presented obvious non neural activity upon visual inspection (e.g., eye blinks, line noise, muscular artefact) were manually rejected. These processing steps were performed using EEGLAB functions (Delorme & Makeig, 2004) for MATLAB. Given that beta CMC is known to occur during steady-state (but not dynamic) phases of force contraction (e.g., Omlor et al., 2007), we focused our analysis of force, EEG and EMG data across the window occurring between 1.5 and 4.5 seconds following the onset stimulus. The initial 1500ms and final 500ms were not considered to minimise data containing initial dynamic contractions or anticipatory relaxations.

### 2.4. Measures

#### Attention allocation

Following the completion of each condition, participants were asked to report how they divided their attention among external factors (e.g., computer screen and force trace), internal factors (e.g., muscular contraction of each finger), and other factors (e.g., mind wandering). Specifically, participants were asked to indicate the percentage (%) of their attention dedicated to each factor, ensuring that the three factors combined to make a total of 100%. These measures were included to elucidate the extent to which our manipulations shifted the balance between external and internal factors for our task as intended.

#### Force control

Force was analyzed to determine the steadiness of each contraction and the extent to which participants accurately maintained their force within their target boundary. Steadiness was defined as the coefficient of variance (CoV), calculated as the ratio between standard deviation and mean during the steady phase. Task accuracy was defined as the percentage of time that participants were able to maintain their force within the target boundary (i.e., 15 ± 5 % MVC).

#### EMG activity

Muscular activity was calculated as the root-mean-square (RMS) of the EMG signal occurring across the steady phase (i.e., 1.5 to 4.5 seconds post stimulus) for each trial. Data were normalised across participants by dividing trial-level RMS by the largest RMS value recorded by each participant across all experimental trials (i.e., % of max RMS).

#### EEG power

Time-frequency decomposition was performed through a short-time Fast Fourier Transform (FFT) conducted on 129 overlapping windows, each of 0.5 s length, with central points ranging from -1.75 to 5.75 s relative to the onset of the auditory go stimulus. Prior to FFT, data points in each window were Hanning tapered and zero padded to reach 1 s. This procedure generated complex-valued coefficients in the time-frequency plane with a precision of 58.6 ms and 1 Hz. EEG power (µV^2^) was calculated for the theta (4 - 6 Hz), alpha (8 – 12 Hz) and beta (15 – 30 Hz) frequency bands as the squared amplitude of each FFT coefficient, which was then averaged across the 51 overlapping segments spanning the steady phase.

#### Corticomuscular coherence

Coherence between the EEG and EMG signals were calculated by magnitude-squared coherence using the following equation:

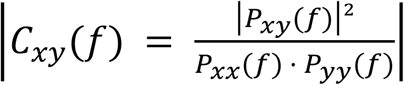

where *P*_*xx*_ and *P*_*yy*_ are the averaged power spectral densities respectively of the EEG and EMG signals throughout the 51 overlapping segments for a given frequency *f*, and *P*_*xy*_(*f*) is the averaged cross-spectral density between the two parameters throughout the segments. The coherence function provides a normative measure of linear correlation on a scale from 0 to 1, where 1 indicates a perfect linear correlation. For multiple corrections across the 16 frequency points within the beta band (i.e., between 15 and 30 Hz), we applied a Bonferroni correction to the following equation defining the significance level of coherence (SL; Rosenberg et al., 1989)

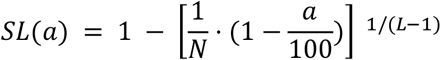

where *a* is the confidence limit (%), *N* is the number of frequency bins, and *L* is the number of overlapping segments. As an *N* of 16, *L* of 51, and *a* of 95 were chosen, the SL was determined to be 0.109 in the present study. This revision eliminates the potential risk that the coherence value is judged to be significant owing to statistical error. We measured the area under the coherence curve and above the significance level in beta frequency range. In this type of area analysis, it is reasonable to take the area under the curve of the coherence because it is a measure of the mean coherence within the considered frequency band (Chakarov et al., 2009; Mendez-Balbuena et al., 2012; Omlor et al., 2007; Trenado et al., 2014). This procedure addresses the overall effect within frequency bands on interest and considers potential shifts in the location of peak coherence across trials, meaning that essential information on the correlated activity between EEG and EMG activity does not get overlooked. Power spectra density plots for all participants are available in the supplementary data to highlight the presence of beta peaks in both EEG and EMG data.

### 2.5. Statistical analysis

The Gaussian distribution of data were checked via Shapiro-Wilk. Consequently, all subsequent analyses were conducted using non-parametric Wilcoxon signed ranks tests to compare our dependent variables across each experimental condition (i.e., external vs internal). Descriptive data are presented by the median and interquartile range. Regional effects of alpha power and beta CMC were analysed through nonparametric permutation testing. The multiple comparison problem (i.e., one test for each channel) was solved through the “maximum statistic” method (M. X. Cohen, 2014) applied to the channel dimension. Namely, we compared the observed Wilcoxon signed ranks *Z* values of each channel between the external and internal conditions with an empirical distribution of *Z* values constructed in the following way. First, we permuted the data by randomly swapping the condition labels (external / internal) within each participant before running a Wilcoxon signed ranks test separately for each channel. We then pooled the *Z* values across channels and stored the two most extreme values (i.e., minimum and maximum). We then repeated this procedure 2,000 times to create a distribution of 4,000 minimum and maximum *Z* values. Finally, we compared the observed (i.e., non-permuted) *Z* values of each channel-wise comparison with the empirical distribution of *Z* values described above, and computed *p* values as the proportion of the permutation *Z* values that were more extreme than the *Z* values of each channel. For alpha power, all 63 channels were included to investigate broad regional effects. For beta CMC, we included the 9 channels spanning the contralateral (left) sensorimotor cortex. Effect sizes for Wilcoxon signed ranks tests were reported as *r*, with values of 0.1, 0.3, and 0.5 reflecting small, medium, and large effects, respectively (Cohen, 1992). Significant differences are presented across figures at the *p* < .05 (*), *p* < .01 (**), and *p* < .001 (***) levels.

## 3. RESULTS

### 3.1. Attention allocation

Participants self-reported a significantly greater allocation of attention towards external factors during the external focus condition (median = 70, iqr = 20%) compared to the internal focus condition (median = 36.11, iqr = 20%), *Z* = 4.090, *p* < .001, *r* = .78. Conversely, participants self-reported a significantly greater allocation of attention towards internal factors during the internal focus condition (median = 55, iqr = 20%) compared to the external focus condition (median = 17, iqr = 18.06%), *Z* = 4.460, *p* < .001, *r* = .85. Attention directed towards other factors was not different between conditions (*p* = .703; Figure 2).

**Figure 2.**
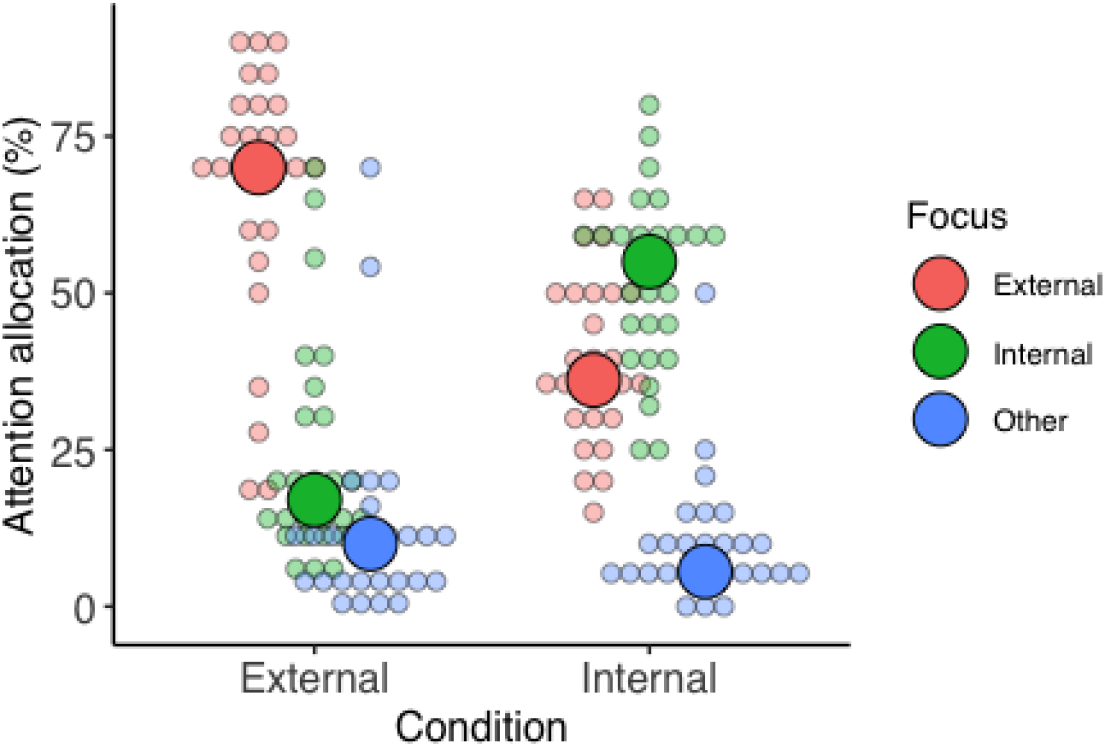
Dot plot displaying the self-reported allocation of attention towards external, internal, and other factors across the experimental conditions. Small points represent individual responses whilst larger points represent the group median.

### 3.2. Force control

A Wilcoxon signed-rank test revealed that force variability (CoV %) was significantly greater during the internal focus condition (median = 3.93, iqr = 1.48%) compared to the external focus condition (median = 2.71, iqr = 0.73%), *Z* = 4.361, *p* < .001, *r* = .84. Additionally, participants were significantly less accurate at the task during the internal focus condition (median = 81.58, iqr = 6%) compared to the external focus condition (median = 91.30, iqr = 10%), *Z* = 4.373, *p* < .001, *r* = .84 (Figure 3).

**Figure 3.**
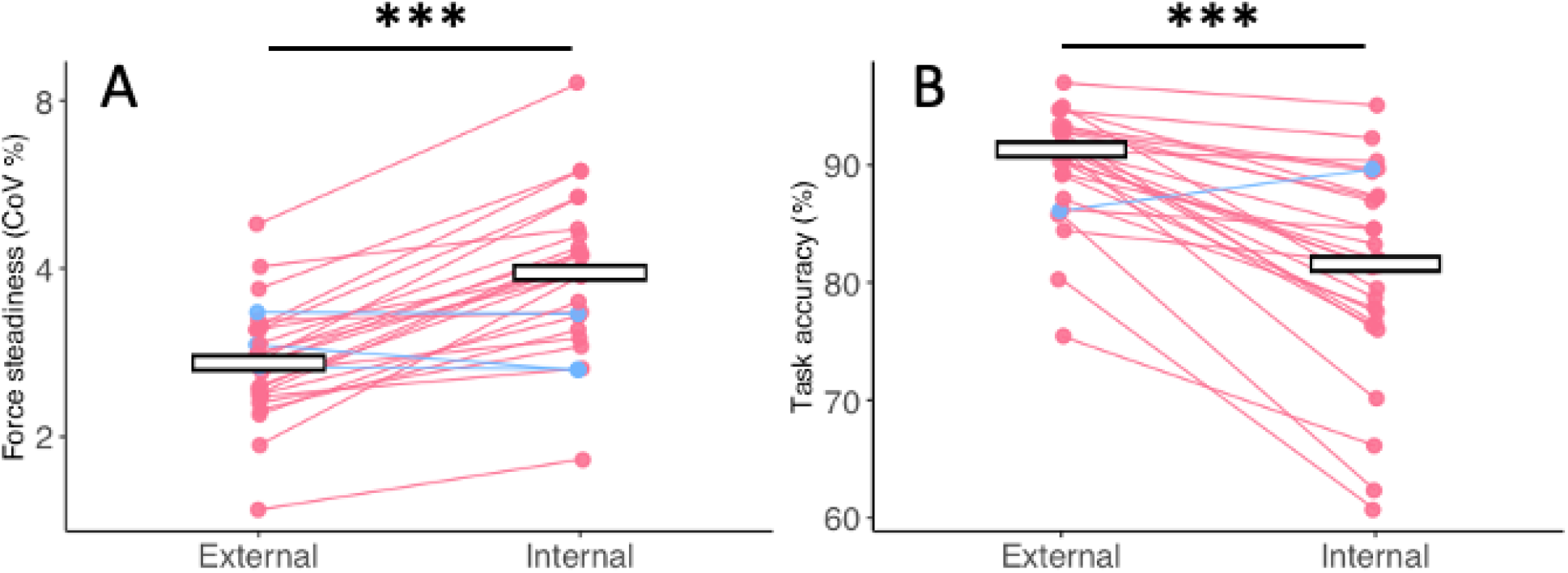
Jitter plots displaying force steadiness (A) and task accuracy (B) across the external and internal conditions. Y-axis values for force steadiness have been log transformed for visualisation purposes. Jitter points represent individual participant means and the horizontal crossbar represents the overall group median. Lines connect jitter points from the same participant across each condition, with a red line representing poorer performance during the internal condition and a blue line representing better performance during the internal condition. Asterisks indicate a significant difference between conditions at the *p* < .001 (***) level.

### 3.3. EMG activity

A Wilcoxon signed ranks test revealed that EMG activity of the flexor carpi radialis was significantly greater during the internal focus condition (median = 59, iqr = 10%) condition compared to the external focus condition (median = 50, iqr = 30%) condition, *Z* = 2.115, *p* = .034, *r* = .41. However, EMG activity of the extensor carpi radialis was not different between conditions, *Z* = 0.361, *p* = .718, *r* = .07 (Figure 4).

**Figure 4.**
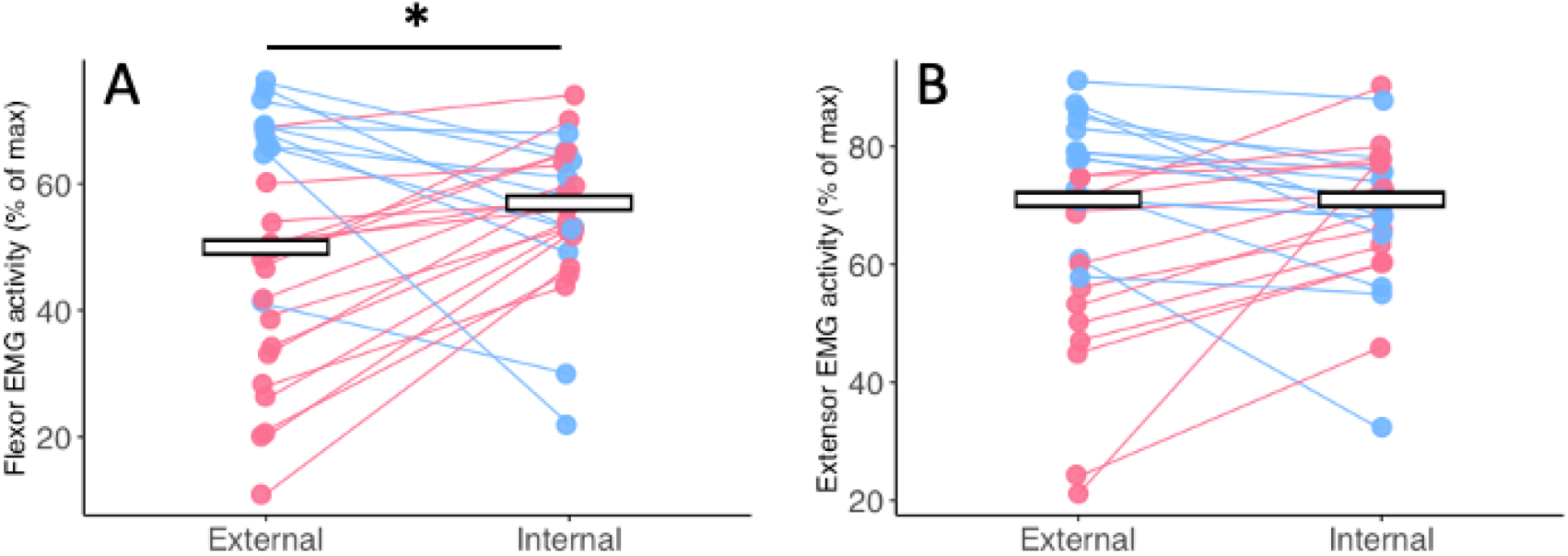
Jitter plots displaying EMG activity (% of max) for the flexor carpi radialis (A) and extensor carpi radialis (B) across the external and internal conditions. Jitter points represent individual participant means and the crossbars represent the overall group median. Lines connect jitter points from the same participant across each condition, with a red line representing increased EMG activity during the internal condition and a blue line representing decreased EMG activity during the internal condition. Asterisks indicate a significant difference between conditions at the *p* < .05 (*) level.

### 3.4. Regional alpha power

Permutation testing to control for the multiple comparisons in the channel x condition dimension identified the observed *Z* values to surpass the critical threshold if they fell below -2.794 or above 2.731. Consequently, results showed that alpha power was significantly greater across a cluster of parieto-occipital channels during the internal focus condition compared to the external focus condition, including O2 (*Z* = 3.003, *p* = .009, *r* = .578), PO4 (Z = 2.931, *p* = .019, *r* = .564), PO6 (*Z* = 3.195, *p* = .003, *r* = .615), PO8 (*Z* = 3.003, *p* = .013, *r* = .578), and Oz (*Z* = 3.051, *p* = .005, *r* = .587; Figure 5).

**Figure 5.**
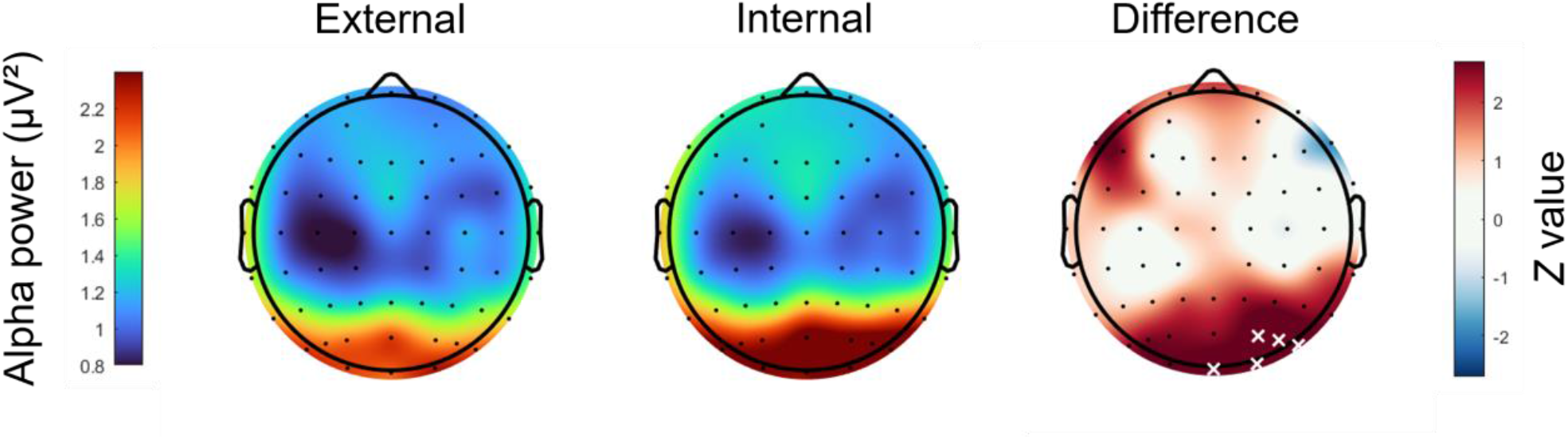
Topographical scalp maps depicting the mean alpha power (μV^2^) recorded during the external focus (left column) and internal focus (middle column) conditions. A scalp map of the paired samples *Z* values comparing alpha power across the external and internal conditions (right column). Areas that are red indicate greater alpha power (i.e., greater inhibition) in the internal condition, whereas areas that are blue indicate lower alpha power in the internal condition. Statistical thresholding was applied using the maximum-statistic permutation testing (Cohen, 2014; Nichols & Holmes, 2002) controlling for multiple comparisons in the channel dimension. Statistically significant channels are indicated by white X markers.

### 3.5. Frontal theta power

A Wilcoxon signed ranks test showed that frontal theta power was significantly greater during the external focus condition (median = 2.11, iqr = 1.65 μV^2^) compared to the internal focus condition (median = 2.00, iqr = 1.52 μV^2^), *Z* = 3.027, *p* = 002, *r* = .583 (Figure 6).

**Figure 6.**
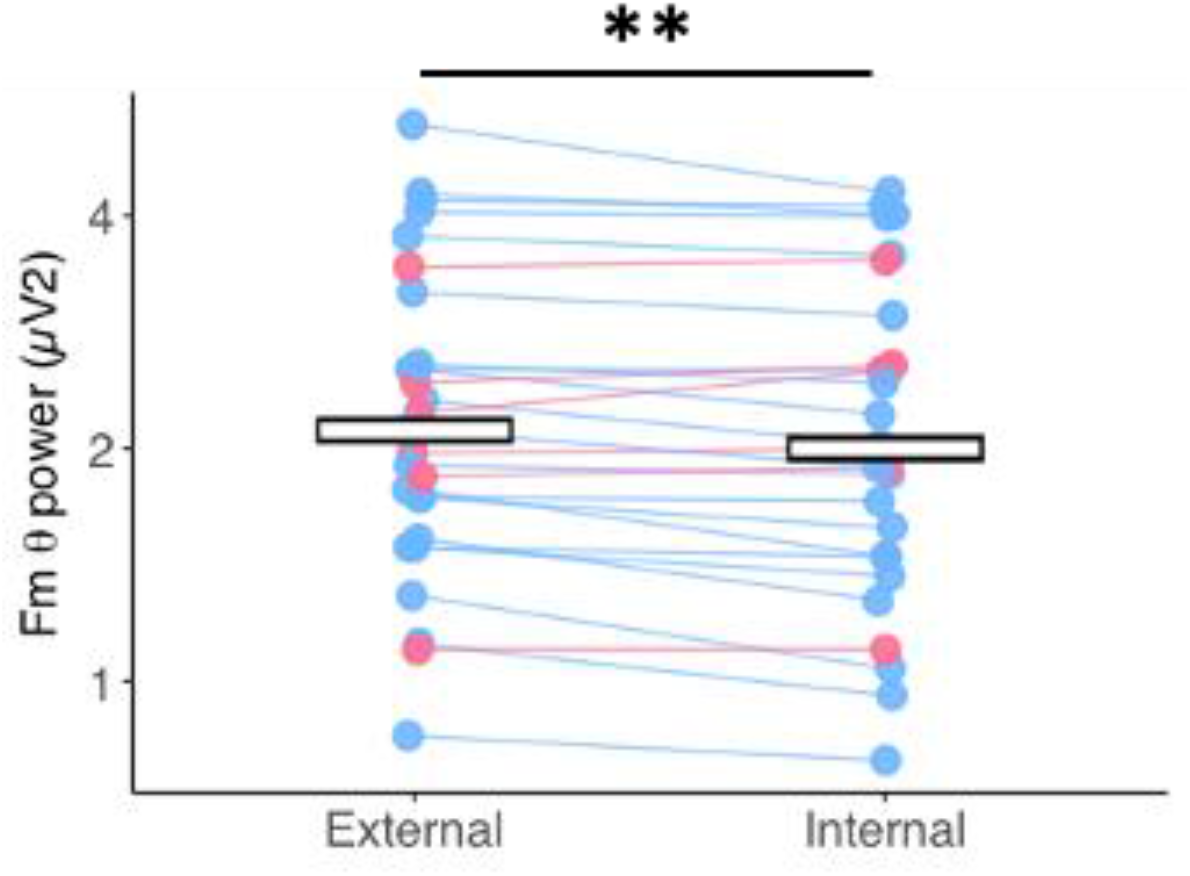
Jitter plots displaying frontal midline theta (Fmθ) power (μV^2^) across the external and internal focus conditions. Jitter points represent individual participant means and the crossbar represents the overall group median. Lines connect jitter points from the same participant across each condition, with red lines represent participants who displayed increased frontal midline theta power during the internal condition and the blue lines represent participants who displayed decreased frontal midline theta power during the internal condition. The y-axis values have been log transformed for visualisation purposes, and asterisks reflect a significant difference between conditions at the *p* < .01 (**) level.

### 3.6. Corticomuscular coherence

#### Flexor carpi radialis

Permutation testing to control for the multiple comparisons in the channel x condition dimension identified the observed *Z* values to surpass the critical threshold if they fell below -2.040 or above 2.040. Consequently, results showed that beta CMC was significantly greater across electrodes CP1 (*Z =* 2.090, *p* = .045, *r* = .402) and C1 (*Z* = 2.427, *p* = .015, *r* = 467) during the external focus condition compared to the internal focus condition (Figure 7).

**Figure 7.**
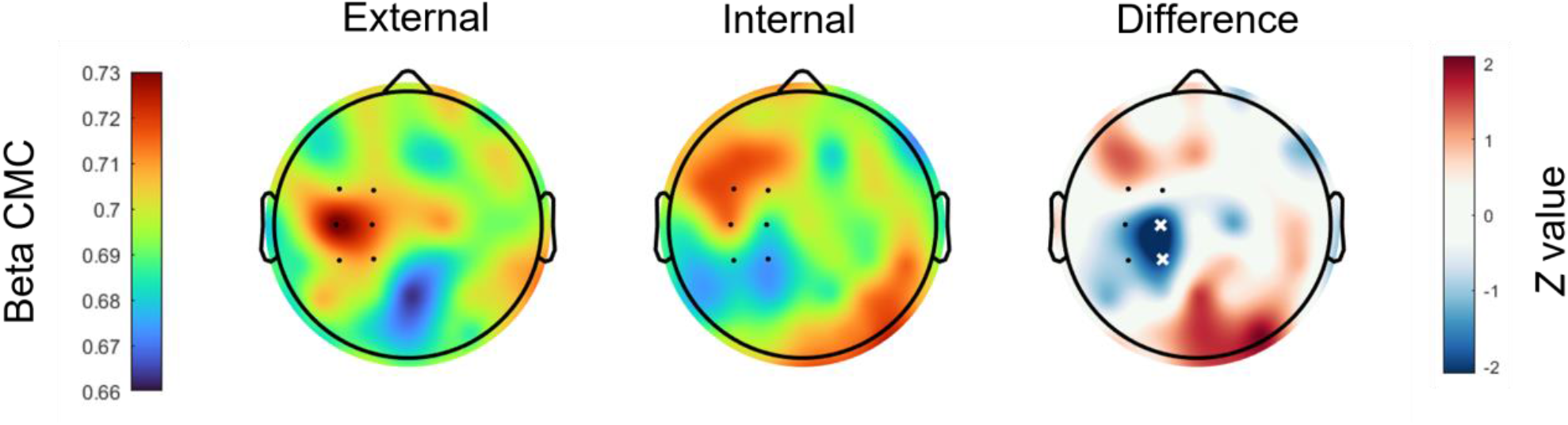
Scalp maps depicting the mean beta CMC recorded during the external (left column) and internal (middle column) focus conditions. Red areas indicate greatest levels of beta CMC whereas blue areas indicate lowest levels of beta CMC. Black dots indicate the channels of the left sensorimotor cortex included in analyses. A scalp map of the paired samples *Z* values comparing beta CMC across the external and internal conditions (right column). Red areas suggest greater beta CMC during the internal focus condition, whereas the blue areas suggest lower beta CMC during the internal focus condition. Statistical thresholding was applied using the maximum-statistic permutation testing (Cohen, 2014; Nichols & Holmes, 2002) controlling for multiple comparisons in the channel dimension. Statistically significant channels are indicated by white X markers.

#### Extensor carpi radialis

Permutation testing to control for the multiple comparisons in the channel x condition dimension identified the observed *Z* values to surpass the critical threshold if they fell below -2.103 or above 2.040. Consequently, no significant differences were observed between the internal and external focus conditions.

## 4. DISCUSSION

This study provided the first cross-modal investigation of how attentional focus alters the kinetic, cortical, muscular, and corticomuscular activity associated with force production in a task requiring precise visuomotor control. As we hypothesised, shifting attentional focus to internal factors (e.g., force across index and middle fingers), as opposed to external factors (e.g., the position of the force trace) impaired force control (accuracy and steadiness) resulting in poorer task performance. The impaired force regulation induced by an internal focus emerged alongside a psychophysiological profile consisting of elevated activity of the forearm muscles, increased alpha activity across the parieto-occipital cortex, decreased frontal midline theta activity, and decreased beta CMC between the forearm and contralateral sensorimotor cortex. The results of this study provide a holistic understanding of how attentional focus strategies alter neuromuscular control.

### 4.1. An internal focus disrupts force regulation and elevates muscular activity

When focusing internally, participants displayed isometric contractions that were less steady (∼1.22% more variable) and less accurate (∼9.72% less accurate) compared to when focusing externally. These findings are synonymous with a large body of literature indicating that an internal focus of attention can broadly disrupt the effectiveness of visuomotor control (Wulf, 2007, 2013). Changes to these task performance metrics were also reflected by differences at the muscular level. When focusing internally participants displayed greater EMG activity of the forearm flexor muscle, suggesting an overflow of muscular activation that was not supportive of optimal task performance. Elevated EMG activity in response to an internal focus is typically interpreted as an attempt to constrain (or “freeze”) the motor system through co-contraction of antagonistic muscles to decrease variability in the movement pattern (Lohse & Sherwood, 2012). However, given that deceased variability would support accuracy in our isometric task (lower CoV), it is unlikely that focusing internally encouraged a “freezing” of action in this instance. Rather, our findings suggest that focusing internally promoted a less efficient muscular output. Interestingly, the forearm flexors specifically contribute to the production of grip force rather than grip force maintenance and relaxation (Ambike et al., 2014). We could, therefore, speculate that the elevated EMG activity of the forearm flexor may not reflect co-contraction per se, but elevated muscular activation in response to a greater frequency and magnitude of large grip force adjustments.

### 4.2. An internal focus elevates parieto-occipital alpha activity

Adopting an internal focus led to greater alpha activity across the parieto-occipital cortex compared to an external focus, aligning with previous findings in word-solving (Ceh et al., 2020) and balance control tasks (Sherman et al., 2021). Our findings therefore extend previous research in the context of a visuomotor force precision task. Alpha activity is widely agreed to reflect regional inhibition, with greater alpha activity indicating increased inhibition and lower alpha activity indicating release from inhibition (Jensen & Mazaheri, 2010; Klimesch et al., 2007). Therefore, our data suggests that elevated posterior alpha activity during an internal focus reflects the functional inhibition of visual processing, likely reflecting the limit of attentional capacity and the consequent need to flexibly divide attentional resources between the different perceptual modalities (visual and proprioceptive) involved with task performance. This is aligned with evidence that attending to visual stimuli whilst ignoring auditory distractors elevates alpha in regions involved with auditory processing whilst simultaneously diminishing alpha in regions involved with visual processing (Mazaheri et al., 2014). Consequently, internally focused attention may induce a state of “*looking without seeing*”, reducing the preparedness to detect and process task-relevant visual information required for the maintenance of task accuracy (Ceh et al., 2020), possibly explaining some of the behavioural differences typically observed between attentional focus strategies.

### 4.3. An internal focus lowers frontal midline theta activity

We also found frontal midline theta activity to be significantly lower during the internal focus condition compared to the external focus condition, corroborating previous findings in balance control (Sherman et al., 2021). Frontal midline theta activity is often proposed to reflect the underlying activity of the anterior cingulate cortex and is commonly measured as an index of the monitoring and updating executive function utilised when exerting top-down volitional attentional control to adjust behaviour in response to feedback error (Cavanagh et al., 2010; Cavanagh & Frank, 2014; Womelsdorf et al., 2010). Heightened frontal midline theta activity has been observed in expert golfers compared to novice golfers when putting (Baumeister et al., 2008) and during the “flow-state” of a mental arithmetic task (Katahira et al., 2018), implying a functional role in maintaining highly focused attention and immersion within a task. Focusing internally therefore appears to disrupt aspects of attentional control that are required for assembling task-relevant information to prime sensory and motor pathways for efficient communication and enhanced task performance.

### 4.4. An internal focus disrupts corticomuscular coherence

Beta CMC between the forearm flexor muscle and EEG channels spanning the contralateral sensorimotor cortex decreased when focusing internally. Whilst the functional interpretation of beta CMC is still debated (e.g., Bourguignon et al., 2019), considerable evidence suggests that the magnitude of CMC is larger for stable compared to unstable contractions (Kristeva et al., 2007; Kristeva-Feige et al., 2002; Witte et al., 2007). Beta CMC is therefore often interpreted to reflect the effective transmission of motor inhibitory beta oscillations from the cortex to the periphery. Decreased CMC can therefore be interpreted as reflecting the departure from a steady-state control of a particular muscle and the disruption of corticomotor inhibition. Our findings therefore align with evidence that focusing internally can disrupt surround inhibition across the motor cortex leading to an overflow of activation at both the cortex and muscle (Kuhn et al., 2017a, 2017b). Interestingly, CMC has been shown to be modulated when applying force to unstable objects that require individuated finger control (Reyes et al., 2017). Decreased beta CMC across the contralateral motor cortex could suggest that focusing internally encouraged individuated finger control and the unbinding of muscles from a synergistic control strategy. Given that our internal instructions specifically focused upon the control of the index and middle fingers, these findings may reflect the decoupling of these fingers from the overall grip force synergy.

### 4.5. Limitations and future directions

It is important to recognise that our findings may not extend beyond the context of our task. To elaborate, our task required high levels of force precision and the upregulation of visual feedback to actively detect and correct for ongoing performance error. As such, it is perhaps unsurprising that inhibition of visual processing during an internal focus might negatively influence task performance. However, research on the preparation for closed-loop aiming tasks has shown that diminished visual processing, as indexed by elevated EEG posterior alpha activity, is associated with better task accuracy (Gallicchio & Ring, 2019). It is therefore possible that disengaging visual processing through internalised attention is a mechanism that disrupts performance only in tasks that contain a dynamic (open-loop) versus stable (closed-loop) visual environment. The possible interpretations of our data are also limited by the absence of a control condition, whereby no attentional focus instructions are given. We chose to exclude a control condition from the design given the large number of trials required for EEG analyses and the possible increase in physical and mental fatigue. Consequently, whilst we interpret these findings as reflecting the negative effects of focusing internally, it is also possible that they represent the beneficial effects of focusing externally. It would therefore be useful to establish how corticomuscular profiles might relate to behavioural (i.e., performance) and perceptual (i.e., attentional strategies) outcomes when no instructions are given during a visuomotor task requiring high levels of precision.

### 4.6. Conclusion

This study provides evidence for a wide range of kinetic and psychophysiological alterations consequent to the adoption of attentional focus strategies in a visuomotor task. We showed that focusing internally impairs the ability to maintain accurate and steady isometric force contractions. These performance deficits were associated with elevated parieto-occipital alpha activity and decreased frontal midline theta activity, suggesting that focusing internally might promote the disengagement of visual processing and disrupt neuronal processes critical to the realisation of cognitive control. An internal focus also increased EMG activity of the forearm flexor and reduced beta CMC between the forearm flexor and the contralateral motor cortex. Our findings build upon evidence that focusing internally can promote an overflow of activation that becomes reciprocated at the cortex and muscles. It is hoped that these findings pave the way for exploring how the well-documented behavioural consequences of attentional strategies can be explained at the corticomuscular level across a wide range of motor tasks and contexts.

## Author contributions

***Johnny Parr***: *conceptualisation, methodology, software, validation, formal analysis, investigation, writing – original draft, visualisation, project administration*. ***Germano Gallicchio***: *conceptualisation, methodology, software, writing – review and editing*. ***Andres Canales-Johnson***: *software, writing – review and editing*. ***Liis Uiga***: *methodology, writing – review and editing*. ***Greg Wood***: *conceptualisation, methodology, investigation, writing – original draft, writing – review and editing*.

## Supplementary material

Supplementary data can be found here: https://osf.io/gktu2/

