## Supplementary figures and images for "Cortical, muscular, and kinetic activity underpinning attentional focus strategies during visuomotor control"

### Supplementary Data

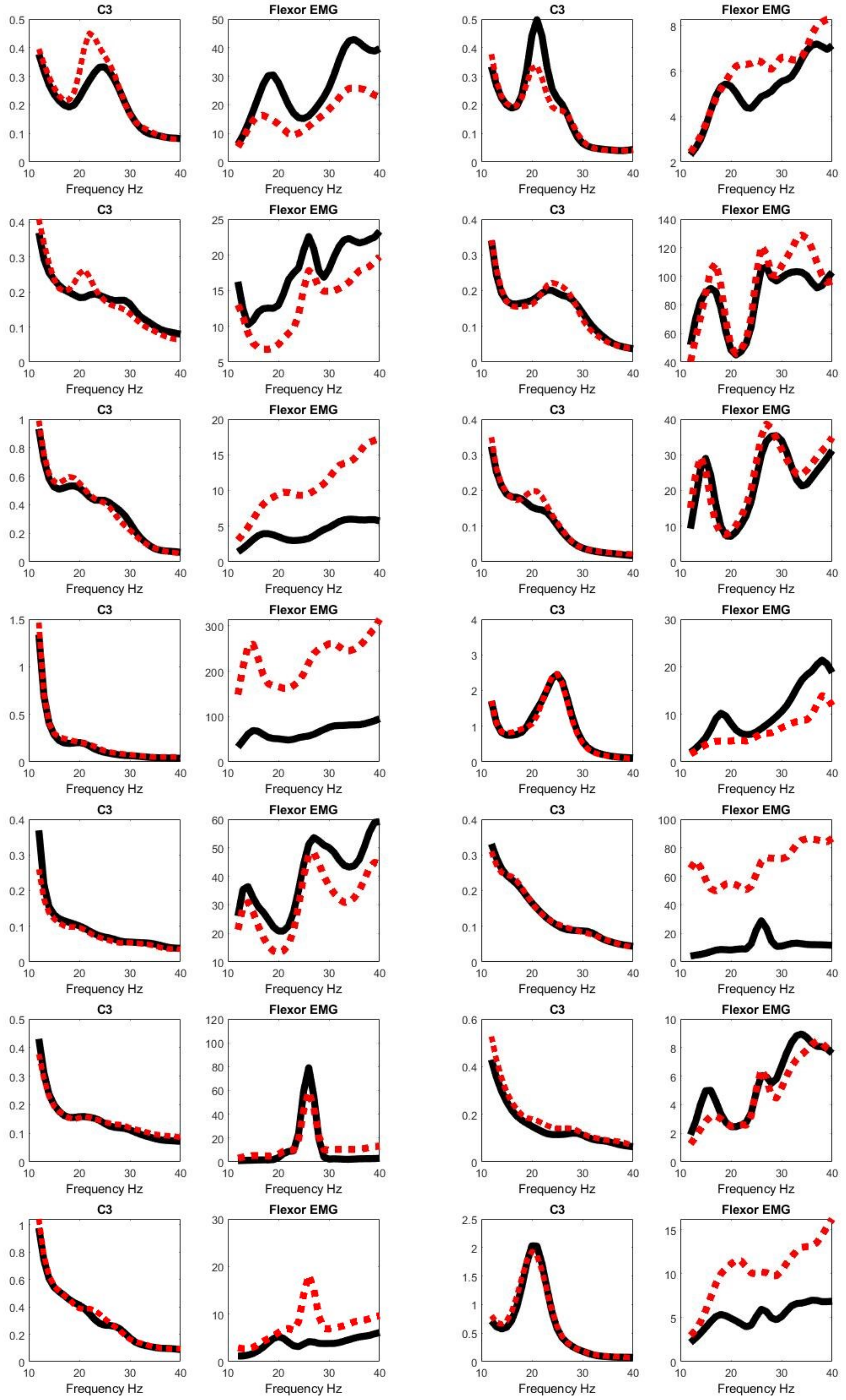

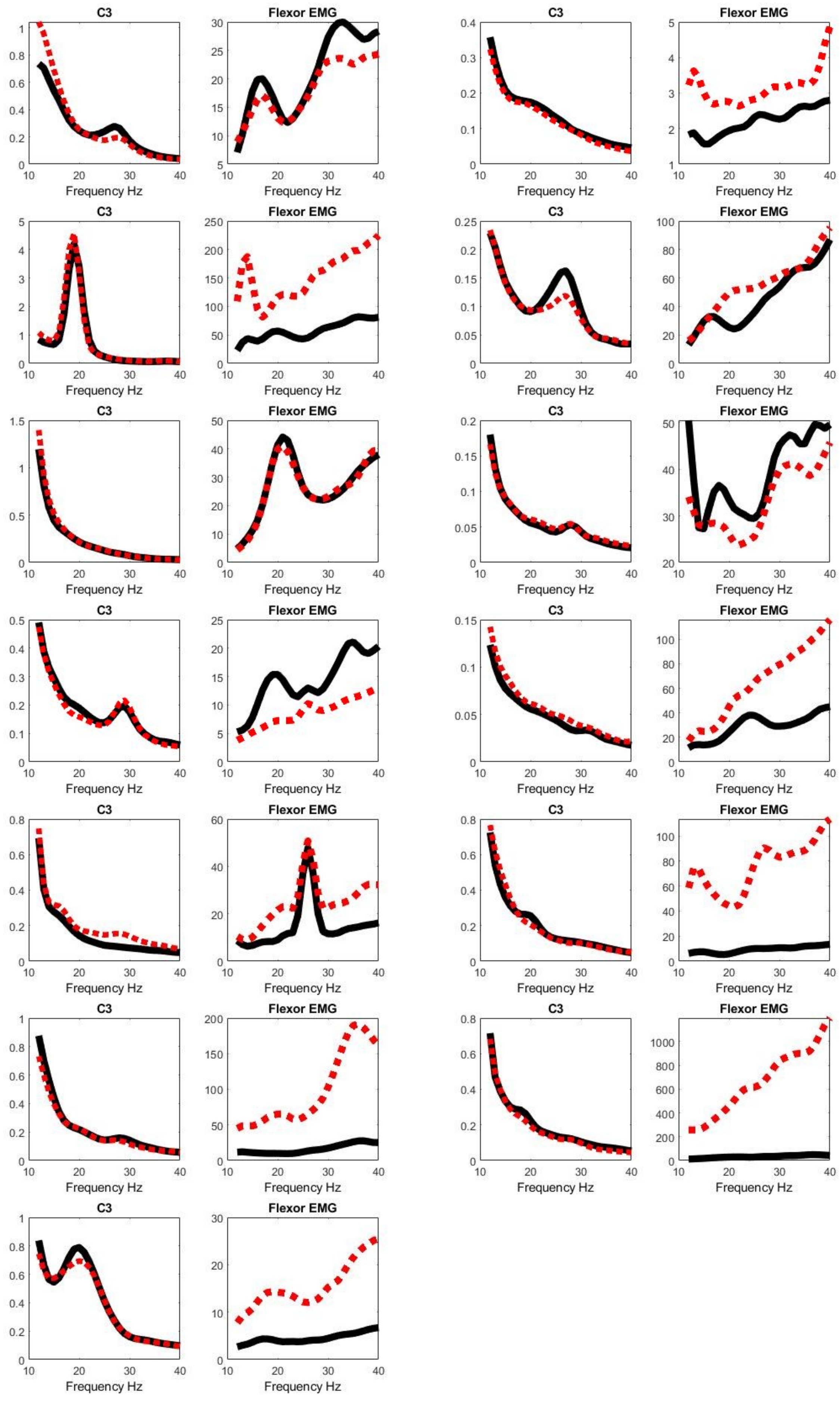
